# High-throughput and automatic structural and developmental root phenotyping on Arabidopsis seedlings

**DOI:** 10.1101/2022.07.13.499903

**Authors:** Romain Fernandez, Amandine Crabos, Morgan Maillard, Philippe Nacry, Christophe Pradal

**Affiliations:** CIRAD, UMR AGAP Institut, F-34398 Montpellier, France; UMR AGAP Institut, Univ Montpellier, CIRAD, INRAE, Institut Agro, F-34398 Montpellier, France; Institute for Plant Sciences of Montpellier (IPSiM), Univ Montpellier, CNRS, INRAE, Institut Agro, Montpellier, France; Inria & LIRMM, Univ Montpellier, CNRS, Montpellier, France

**Keywords:** High-throughput phenotyping, computer vision, dynamic analysis, root system architecture, tracking

## Abstract

**Background:** High-throughput phenotyping is crucial for the genetic and molecular understanding of adaptive root system development. In recent years, imaging automata have been developed to acquire the root system architecture of many genotypes grown in Petri dishes to explore the Genetic x Environment (GxE) interaction. There is now an increasing interest in understanding the dynamics of the adaptive responses, such as the organ apparition or the growth rate. However, due to the increasing complexity of root architectures in development, the accurate description of the topology, geometry, and dynamics of a growing root system remains a challenge.

**Results:** We designed a high-throughput phenotyping method, combining an imaging device and an automatic analysis pipeline based on registration and topological tracking, capable of accurately describing the topology and geometry of observed root systems in 2D+t. The method was tested on a challenging Arabidopsis seedling dataset, including numerous root occlusions and crossovers. Static phenes are estimated with high accuracy (*R*^2^ = 0.996 and 0, 923 for primary and second-order roots length, respectively). These performances are similar to state-of-the-art results obtained on root systems of equal or lower complexity. In addition, our pipeline estimates dynamic phenes accurately between two successive observations (*R*^2^ = 0. 938 for lateral root growth).

**Conclusions:** We designed a novel method of root tracking that accurately and automatically measures both static and dynamic RSA parameters from a novel high-throughput root phenotyping platform. It has been used to characterize developing patterns of root systems grown under various environmental conditions. It provides a solid basis to explore the GxE interaction controlling the dynamics of root system architecture adaptive responses. In future work, our approach will be adapted to a wider range of imaging configurations and species.

## Background

Plants expand their root system to meet their needs for water and nutrients. Root system shape, structure, and plasticity impact its efficiency and condition the plant fitness in fluctuating environments (Balduzzi *et al*., 2017). The root system architecture (RSA) is dramatically affected by spatial and temporal changes in the soil environment that, in turn, will modify its water and nutrients capture efficiency (Boursiac et al., 2022). These adaptive responses, including branching angles and frequencies of lateral roots, rely on growth rate modifications of the different root classes (e.g., primary, seminal, lateral, or adventitious roots). The optimal distribution of roots in a given environment allows plants to forage efficiently for resources. There is a need to explore the link between RSA and its dynamics, plant physiology, and agronomic traits to implement plant breeding programs (Lynch, 2022). Such goals involve designing non-destructive high-throughput phenotyping systems. Non-destructive measurements are indispensable to periodically quantify the phenotype of the same plant all along time and then decipher the dynamics of the adaptive responses. Because roots are the below-ground organs and represent the hidden half of the plant, non-destructive root phenotyping is technically challenging for high throughput. One approach to quantify root growth and geometry under a wide range of growth conditions is to use a transparent medium, such as agarose gels, a common cultivation practice for the model plant Arabidopsis and other species. Several semi-automatic tools have been developed to assist root phenotyping on agar plates (Yasrab *et al*., 2019; Nagel et al., 2020; Gaggion *et al*., 2021; Möller *et al*., 2021; Ohlsson *et al*., 2021). However, these devices are either limited in the number of plates imaged or restricted in temporal resolution. This situation hampers potentially useful phenotypical parameters of individual roots that may be linked to the temporal dynamics of growth. Here we present HIRROS, an automated and non-destructive visualization automat for 200 Petri dishes and an image analysis pipeline allowing high-throughput temporal phenotyping of *seedlings* RSA. to support the massive phene collection required for proper genetic studies in response to fluctuating environments.

Automatic extraction of RSA is a current bottleneck and an active topic in bio-image analysis. The community attempts to develop solutions to process root systems 2D images (Lobet *et al*., 2011; Yasrab *et al*., 2019; Takahashi, 2021), 3D CT observations (Mairhofer et al., 2016; Gaggion *et al*., 2021; Herrero-Huerta et al., 2021), or multi-angle 3D reconstructions (Symonova *et al*., 2015); see (Ndour *et al*. 2017) for a detailed description of standard root system phenotyping setups. The proposed processing pipelines for reconstructing RSA from 2D or 3D images typically include a step of image segmentation (root detection and background removal) and a step of RSA reconstruction by optimal path search in the segmented structures (Yasrab *et al*., 2019; Möller *et al*. 2021). The first step is a pixel classification task; previous works proposed various segmentation techniques, including pixel classifiers controlled by fixed or trainable parameters. Recent works exhibit significant advances using deep convolutional networks (Yasrab *et al*., *2019*, Möller *et al*., 2021: Smith *et al*., 2020). Unfortunately, these methods require vast training data and offer no guarantee when applied to a new dataset with different characteristics (species, observation device, bogus phenotype). Some authors partially addressed this concern using transfer learning (Yasrab *et al*., *2019*), allowing the end-user to transfer the method to a new dataset after re-training the network.

The second step, aiming at RSA reconstruction, relies on algorithms extracting root paths from segmented structures, interpreting pixel sets as connected root systems to output higher-level primitives, i.e., plant organs and connections. Biologists need to assess that this reconstruction process solves the architecture at the organ scale. This feature is required to distinguish local responses from systemic responses (elongation speed, branching density) (Ruffel et al., 2011; Rellan-Alvarez et al., 2016; Maurel et al., 2020), but offering such guarantees remains a challenge. There are several difficulties in current solutions, mainly due to the capacity to deal with root intersections and root contacts (Symonova *et al*., 2015, Gaggion *et al*., 2021; Yasrab *et al*., *2019*). Each new crossing doubles the number of possible root paths, which causes a combinatorial explosion of the reconstruction problem. Some authors admitted that the root crossing issue is a significant failure cause without proposing any solution to tackle it (Symonova *et al*., 2015). A global optimization algorithm could provide an optimal solution to this problem. However, the most popular approach in the literature is reconstructing the different organs sequentially (Yasrab *et al*., *2019)*. This method is more or less equivalent to addressing the root crossing problem with a greedy algorithm, leading to local optima.

With the development of high-throughput time-lapse observations, there is a shift to spatiotemporal data, adding another layer of complexity. The most common way to achieve spatiotemporal architecture reconstruction is to consider time points as separated data. These time points are processed using the conventional two-step pipeline before exploring the correspondences between the extracted architectures over time. This operation requires advanced matching methods (Boudon *et al*., 2014; Symonova et al., 2015; Gaggion et al., 2021) and extra processing steps that could induce a performance drop when cumulating the errors of the separate reconstructions. Some authors attempted to integrate temporal information in the reconstruction process to address this issue. Gaggion et al. (2021) sidestep the geometric alignment task by fixing the specimen on the imaging device during the time course. With this setup guaranteeing a proper alignment of the successive observations, the authors enhance the temporal consistency of the obtained segmentation using a weighted average over the time series. The drawback of this method is the need to provide N observation devices for the N specimen (or for the N petri dishes, eventually), which is incompatible with high-throughput experiments. Finally, Yasrab *et al*. (2021) proposed a novel perspective on the whole problem, training a Future-GAN network to reproduce the developmental behavior of the root systems. This method showed high performance for geometry reconstruction but is prone to errors in topology reconstruction.

RSA complexity increases progressively in time series, making the time information a lever for solving the architecture problem: space and time can be combined to discriminate organ connections from root crossing. In this work, we propose a novel method for the automatic reconstruction of RSA integrating spatiotemporal information as a whole in an algorithm processing 2D+t data. First, a registration pipeline is built to address the alignment issue. Then, root systems are segmented and labeled with root apparition time. A novel root topological tracking algorithm is applied that combines temporal and spatial information and solves ambiguities introduced by root crossing. This pipeline is applied on 1000 time-lapse images of Arabidopsis roots grown in petri dishes (N=1000 plants observed during 21 timesteps, timestep=8 hours), and the result is saved in the RSML format (Lobet *et al*., 2015), enabling the extraction of new spatiotemporal phenes. Our automatic pipeline, evaluated over N=200 plants imaged and expertized over 21 timesteps, shows state-of-the-art performances on high-throughput data. Finally, the development of architectures is reconstructed from time-lapse, providing unseen insights into root system growth.

## Methods

### HIRROS: automated platform for RSA imaging

The HIgh Resolution ROot Scanner (HIRROS) setup is a device for automated and non-destructive visualization of root growth and architecture of Arabidopsis thaliana plants or small seedlings (Fig. 1). The imaging automat is located in a dedicated growth chamber allowing temperature, hygrometry, photoperiod, and light intensity to be adjusted according to the user’s requirements. (Root phenotyping platform webpage: [Nacry, 2022]).

**Figure 1:**
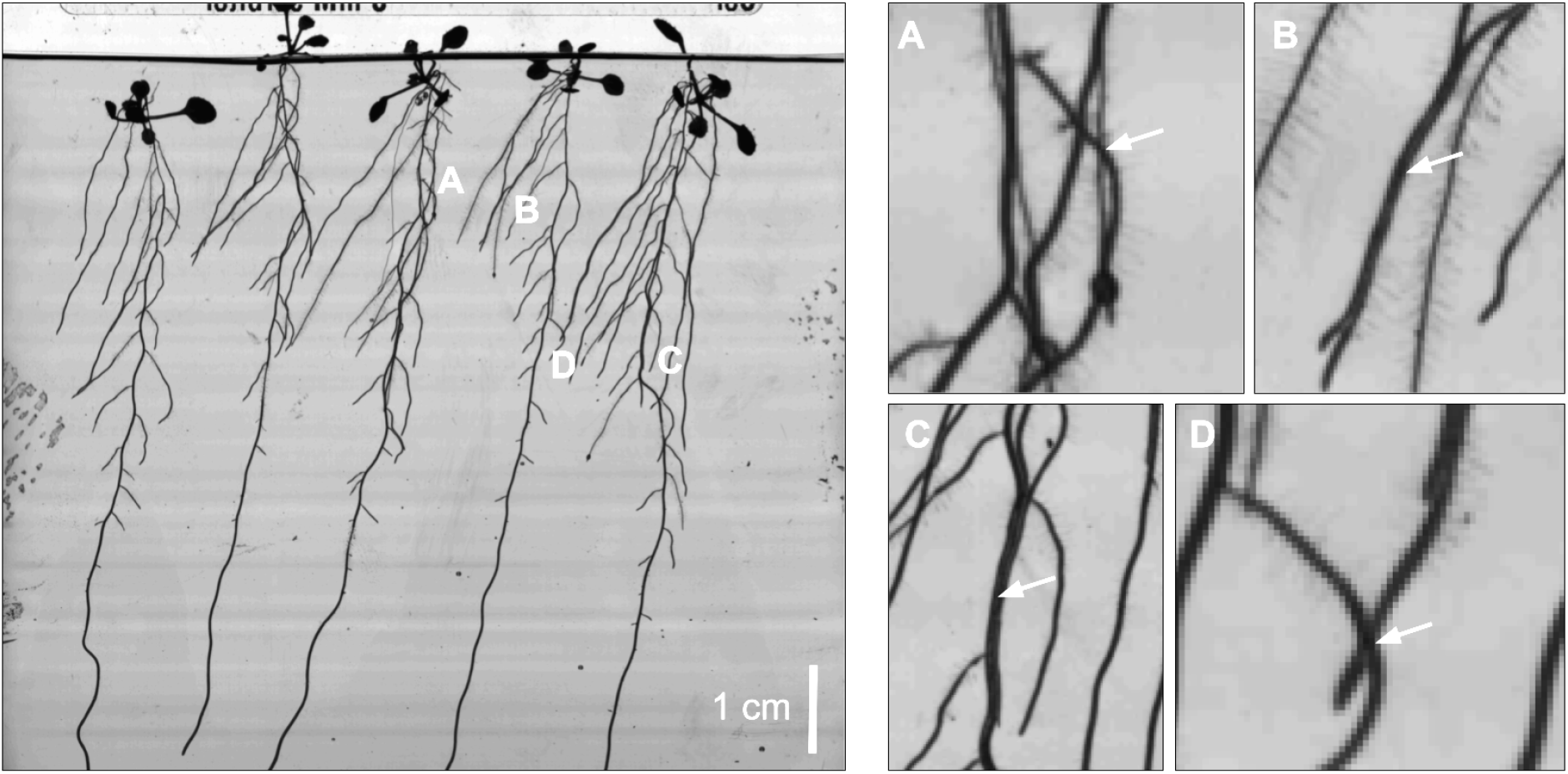
Challenges of root system architecture reconstruction. Left, an image of Arabidopsis seedlings with a developed architecture, observed in a petri dish. (A) When observed in 2D, these root systems exhibit numerous root crossings, which induce possible root tracking errors, and can even fool experts when annotating static images without a “temporal insight.” In addition, multiple roots can follow each other for an arbitrarily long time (B), making them hard to disentangle. Finally, second-order roots sometimes hide below the primary root (C), and roots from two adjacent plants could cross (D). These points are severe issues for static reconstruction algorithms and greedy reconstruction processes, requiring addressing this problem with a 2D+t global algorithm.

HIRROS is designed to use either 200 small (120 × 120 mm) or 72 large (245 × 245 mm) square Petri dishes, which are filled with agar medium and placed upright in holders to allow root growth along a vertical plane at the surface of the culture medium. Once the medium is solidified, a one-centimeter wide strip of medium is removed to avoid contact of cotyledons and leaves with the culture medium and maintain shoots vertically (Fig. 1). Plate lids are wiped with tween20 to reduce condensation and water droplet formation. Five pre-germinated seedlings are evenly distributed, and the plates are sealed with four pieces of tape to allow gas exchanges. The Petri dishes are then placed into holders to maintain them vertically and make them transportable within the automated HIRROS setup. Holder motions are driven by worms allowing lateral and forward movements. The imaging system that is protected from parasitic light is located beside the culture zone (Fig. 1). Plates are transferred to the imaging cart. For illumination, the plates are backlighted with a white collimated LED (Fig. 1). This allows a high contrast (even for thin and almost translucent Arabidopsis roots) and reduces reflection on the plate’s lid. Plates are imaged using a 16MP linear camera coupled to a telecentric lens located at 50 cm of the plate (Fig. 1). This optical setup increases the depth of field. It suppresses image distortions allowing imaging of small and large plates without adjustment. It has a resolution of 19 μm/px (image size 8 MB, TIFF for small plates and 32 MB for large plates). Metadata describing the experiment associated with each image and stored locally in a folder labeled with the experiment information and the imaging date. Illumination is switched on and off automatically and synchronized with the camera during image acquisition. Once imaged, the plate moves back to the holder that moves one position further to place the next Petri dish in the row in front of the imaging cart until all plates are recorded. The holder moves back to the culture zone, and the next holder enters the imaging process. The imaging frequency can be adjusted between 2 and 24 hours, and up to 1000 plants can be imaged in less than 80 minutes.

### Image processing pipeline

Our processing pipeline (see figure 1) is fully automatic. It comprises four steps (see figure 1): 1) time series registration, 2) 2D+t segmentation, 3) root topological tracking, and 4) architecture reconstruction.

#### 1 Time series registration

The pipeline’s first operation corrects the root system’s misalignment in the 2D+t image sequence 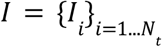, induced by the displacement of the plate in the imaging automat. We identify two sources of misalignment: i) movements of the box relative to the imaging device and ii) deformations of the root system in the culture medium induced by plate displacement. We correct these distortions using a registration pipeline developed with the Fijiyama plugin libraries (Fernandez *et al*., 2021).

We fix the geometry of the last image 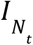, considered the reference image because it contains all the structures present in the previous images. We determine (*N*_*t*_ − 1) geometric transformations {*T*_*t*_} to align the other images with the reference image 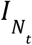,. Each estimated transformation *T*_*t*_ comprises a rigid transformation (translation + rotation), and a non-linear transformation (dense vector field) estimated successively. In this setup, we assume that the rigid transformation is required to correct the movement of the plate, while the non-linear transformation is required to correct the slight deformations of the root system in the culture medium. These transformations are computed from local correspondences using the blockmatching algorithm (Ourselin *et al*., 2000), integrated into a daisy-chain strategy: each image is registered with the next image in the sequence. Then the successive transformations are composed and applied at once for each image.

#### 2 Time series segmentation

The image time series shows the roots growing over time. The difference between two successive images highlights the newly appeared segments of the roots. By working on these differences over the image sequence, one can distinguish the roots from the image’s background, thus segmenting roots and associating each root segment and apparition time. Thus, we define the two purposes of this segmentation step: i) identify which image pixels are traversed by the root systems during the sequence (binary segmentation of RSA), and ii) estimate the corresponding traversal times.

In our dataset, background pixels have medium intensity while root pixels have low intensity. However, segmenting roots by thresholding each image separately is not an option, as other structures (box border, condensation, dirtiness) may have similar characteristics to roots, and pixels intensity varies between the root central line (low intensities) and root border (medium intensities).

Nevertheless, the registered image sequence allows studying the time sequence of intensities of each pixel and detecting in this sequence the event associated with the detection of a root (i.e., when a root reaches this pixel). The time sequence of intensity of a pixel traversed by a root begins with high values corresponding to the background, then undergoes a sudden decrease of intensity at the moment when the root reaches this pixel, and finally finishes with low values corresponding to root tissues. We process each pixel individually and identify this pattern by computing the maximum mean shift over its time series of intensity. We identify as “root pixels” those whose mean shift value is greater than a fixed threshold *s*_1_ = 25 (given that the average intensity difference measured between the centerlines of the roots and the background is approximately 100). Among these pixels, we compute the difference between successive values in the time sequence and identify the observation time following the maximal intensity drop. If this maximum intensity drop is lower than a fixed threshold *s*_2_ = 10, the pixel is rejected and identified as background. Else, the index of the maximal intensity drop is identified as the “apparition time” of the root at this given pixel location. The pixels selected by these two thresholding operations constitute a binary segmentation of the root systems. Each selected pixel is given a label defining the root apparition time (see Figure 4).

We run a topological and geometrical characterisation of the binary segmentation of the root systems to eliminate various outliers: nightly condensation and dirtiness falling on the dish during the experiment. We compute the 4-connected components and reject the pixels of components smaller than 2000 pixels. In our setup, this threshold is barely equivalent to rejecting any candidate root system in which cumulative organ length at the last observation time point is lower than 38 mm, which is suited to our experiment.

This segmentation step results in a 2D labeled image *S* (see figure 2), whose pixels *p* are given a label *S*(*p*) between 0 and *N*_*t*_ ; the label “0” identifies non-root pixels, and the other possible labels identify the index of the first image where the root is present.

**Figure 2:**
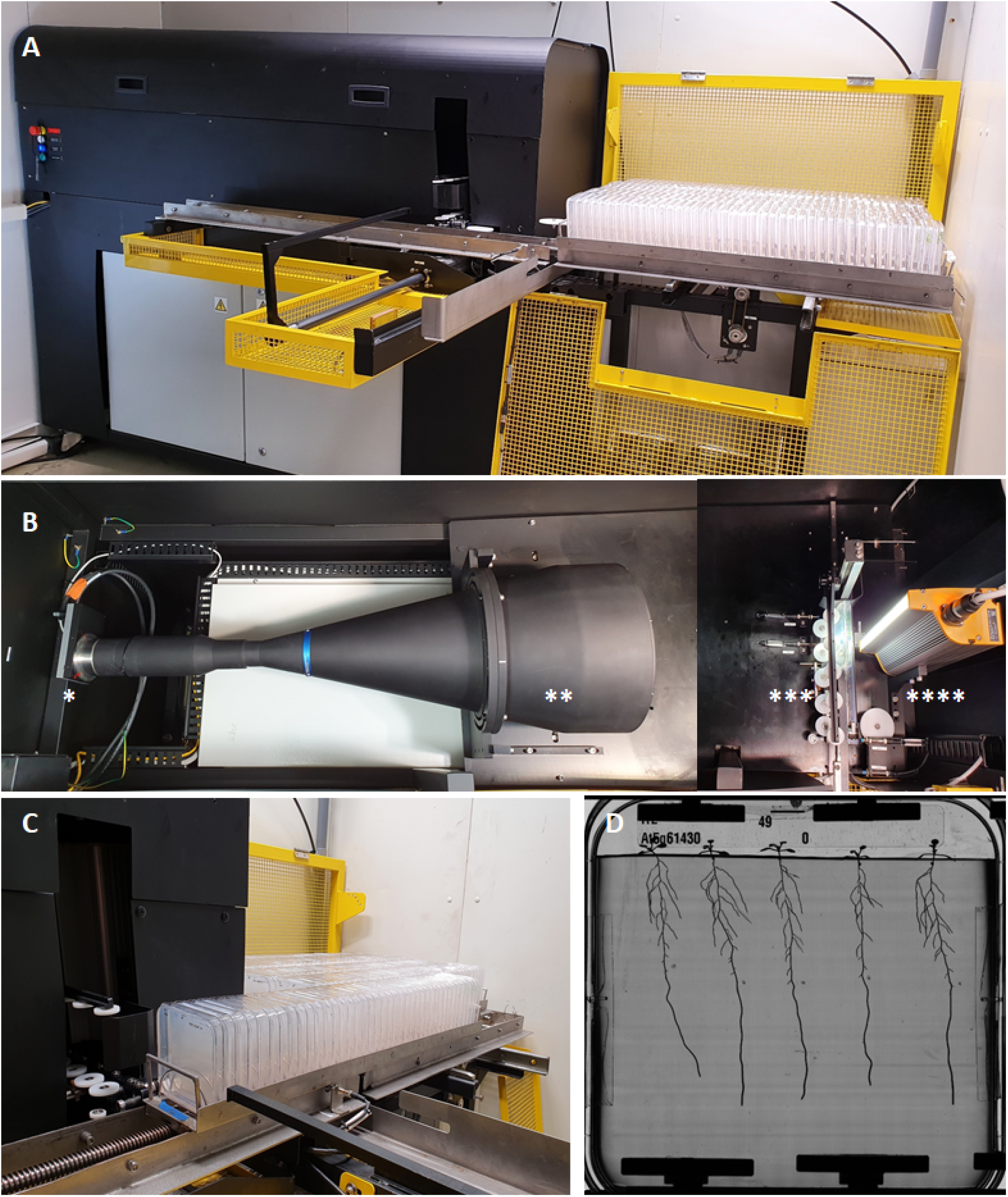
The HIgh Resolution ROot Scanner (HIRROS) setup for automated and non-destructive visualization of root architecture of seedlings grown in agar plates. Global view of the device (A) with the culture zone on the right side and the imaging zone on the left side. Overview of the imaging zone (B), which is composed of a 16Mpixel linear camera (*), a telecentric objective of a diameter of 25cm (**), a mobile imaging cart (***) holding plates during imaging, and a collimated LED backlight (****). The square Petri dishes are maintained vertically in metal racks. They are stored in the culture zone (right side of the device). The rack moves to the imaging area (C) for image acquisition, and the plate is pushed in the imaging with a moving arm. Once the image is captured, the plate moves back to its initial location, and the rack moves to image the next plate. Once all plates are imaged, the rack moves back to the culture zone to proceed to the next one. Representative original grayscale image of Arabidopsis seedlings taken by the camera (D). 200 Petri dishes of 12×12cm or 72 Petri dishes of 24×24 cm containing up to 1000 Arabidopsis plants fit into HIRROS.

#### 3 Root topological tracking

Multi-particle tracking can be challenging, especially if the number of particles is not constant, which requires managing the creation and destruction of tracks in time (Luo et al., 2021). An additional difficulty comes when the temporal sampling is sparse with respect to the particle velocity, inducing possible confusion between crossing tracks or neighbouring tracks. For the tracking of roots in the time series, we propose a solution based on a global tracking algorithm working on the full-time series. In the previous step, the pipeline outputs the growing architecture as a fixed architecture with time-stamped elements. During root topological tracking, the method tracks structures by using these timestamps. In steps 3-a, the algorithm computes a directed Region Adjacency Graph equivalent to the segmentation. In 3-b, it extracts a covering forest of minimal weight from it, and provides the skeleton of the primary and secondary roots. Then in 3-c, it processes the root crossings by a global optimization algorithm (see figure 4).

#### 3-a) Representation of roots growth space using a weighted directed region adjacency graph

After registration, we expect that the root system appears in the sequence of images as a sequence of Russian dolls: assuming that a root does not disappear (by dying or being removed) if we note *P*_*t*_ the set of pixels covered by the roots at the time *t, P*_*t*_ ⊆ *P*_*t*+*k*_ (*k* > 0), and the difference *P*_*t*+*k*_ \ *P*_*t*_ is the additional surface of root system generated by root tips growth between *t* and *t* + *k*. Thus, we assume that a connected component of pixels in *S* sharing the same label *t* is an additional surface covered between the observation time number *t* and *t* + 1 by a part of the root systems. With this assumption, root trajectories can be tracked by identifying the optimal pixel paths in *S*, beginning with pixels labeled 1 (sequence start) and finishing with the label *N*_*t*_ (last observation time point). We formalize a global solution to the tracking problem by searching a minimum cost spanning forest in the weighted directed region adjacency graph *G* = (*V*_*G*_, *E*_*G*_) of the labeled image *S* (see figure 3). We build *G* as follows:

**Figure 3:**
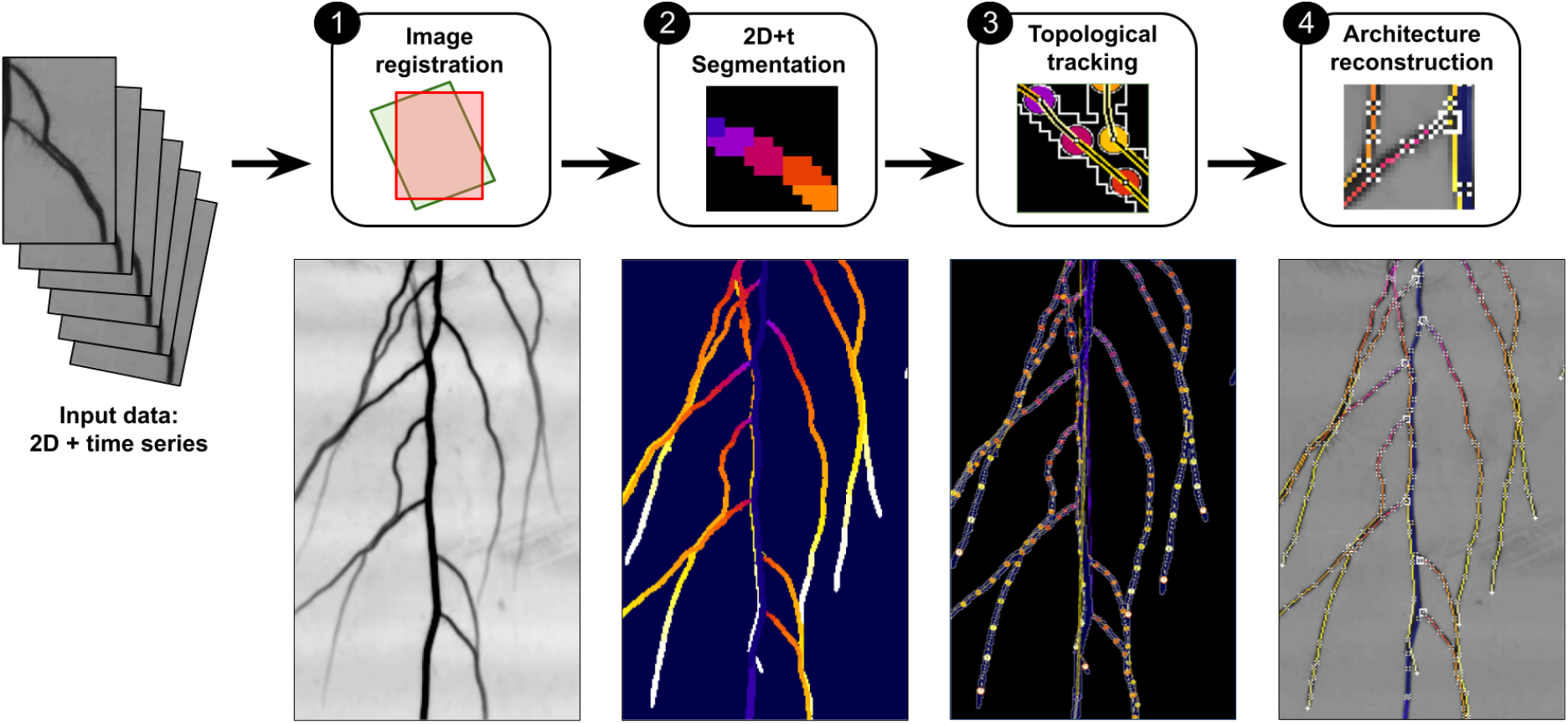
Processing pipeline overview. Upper line: pipeline steps, bottom line: output of the successive steps.

- Each 8-connected component of pixels *p* ∈ *S* sharing the same label *t*(*p*) in *S* is represented by a vertex *𝑣* ∈ *V*_*G*_ with label *t*(*𝑣*) = *t*(*p*).
- Two vertices *𝑣* _1_, *𝑣* _2_ are connected with a (directed) edge *e* = (*𝑣*_1_, *𝑣*_2_) if and only if *t*(*𝑣*_1_) < *t*(*𝑣*_2_) (edges are directed according to forward time direction) and the two corresponding 8-connected components in *S* are 4-neighbors (they share at least a pixel edge).

The weight *p*(*a*) of an edge *e* = (*𝑣*_1_, *𝑣*_2_) is defined by:

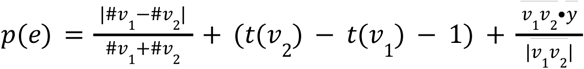

with

- #*𝑣* the number of pixels contained in the corresponding connected component in *S*.
- *t*(*𝑣*) the label of *𝑣*.
- 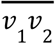 is the vector from *𝑣*_1_ to *𝑣*_2_ and 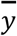 is the Y-axis normalized vector.

The weight *p*(*e*) of an edge is small (the edge is more “likely” to belong to an actual root trajectory) if i) the edge connects two vertices which have a comparable surface, ii) the two vertices labels (structures apparition time) are close, and iii) the vector connecting the target from the source is directed downwards, which is the preferred orientation of root growth.

#### 3-b) Extraction of root system skeleton

At this step, we extract the root systems skeleton in the graph by identifying the paths corresponding to primary roots and lateral roots without considering lateral root crossings. We describe this reconstruction algorithm for architecture composed of primary roots and lateral roots (second-order organs inserted on the primary), which is suited to arabidopsis seedlings.

### Primary roots

The primary root of seedlings is the first organ of the root system and tends to be larger than the lateral roots, have a higher growth rate and point downwards. Consequently, the corresponding path in the graph is composed of edges of minimum weight p(e). At the beginning of the experiment, we assume that the primary axes of the N plants (Np=5 plants per box in our setup) have no branching. We then identify the corresponding vertices in G, which are the Np vertices associated with the largest connected component of timestamp 1 in the segmentation S. From each of such vertices, we extract the path of the primary axis, which is the longest greedy min-cost path. Vertices and edges traversed in this step are labeled primary, and all other edges and vertices in G are labeled lateral. In agreement with the root hierarchy, all lateral edges incident to a primary vertex are invalid and eliminated from the graph. Finally, the algorithm filters out outliers by computing an extraction of connected components in G from the primary vertices and rejecting all vertices not included in the final components.

### Lateral roots

The algorithm identifies the path corresponding to lateral roots by selecting in the graph the vertices and edges that are likely to correspond to actual lateral root paths (see figure 3). This action of classifying between “relevant” and “irrelevant” edges is an action of graph pruning that we run in two steps. First, we aim to extract a subpart of the graph whose topology corresponds to that of a set of root systems, which is a forest structure. To that end, we extract a forest from the graph by computing the minimal directed spanning forest *F* = (*V*_*F*_, *E*_*F*_) using the Edmonds algorithm (Edmonds, J., 1967). In *F*, each vertex has a single incoming arc. Second, we try to ensure that each path corresponding to a lateral root has no branching, according to our dataset composed of arabidopsis seedlings. In order to avoid the branching of lateral roots, we check each lateral vertex and keep only their min-cost outgoing arc.

### 3-c) Resolution of root crossing ambiguities

By pruning the graph, parts of actual root paths corresponding to root crossings could be disconnected. This point is investigated and processed in this step using a global algorithm. If a root *r*_2_ crossed the path of another root *r*_1_ during the time sequence, assuming that *r*_2_ traversed the area at least one timestep after *r*_1_, the region adjacency graph in 3-a) shows a typical “X” pattern, where the path of *r*_2_ cross at least one edge in the wrong direction. This pattern leads to an upstream disconnection of the *r*_2_ path and a downstream disconnection of the *r*_2_ path during graph pruning in 3-b), which lets an additional root start and root end in the graph (see figure 3). To process these cases using a global algorithm, we first identify the two following vertices sets in *F* : the “root starts” *V*_*start*_, including every lateral vertex with no incoming arc, and the “root stops” *V*_*stop*_, including every vertex with no outgoing arc. As these elements are possibly parts of disconnected pathways, we target to identify the best pairwise matches (root stops with root starts) in order to reconnect them. We define a connection cost function to compute the optimal set of pairwise matches between elements of these two sets. The cost function is defined to indicate the likelihood of the connection, computed as the weighted average of multiple geometrical and topological continuity features, replaced by majoring values when the elements to connect are not long enough to provide the needed information. The features used for the cost function are the edge weight function *p*(*a*) (see 3-a), the angle between root start and root end, the relative speed difference between the root start and the root stop, and a topological score. The topological score is computed by evaluating the minimal path between the root end and root start in the original region adjacency graph, evaluating the number of edges crossed in the right or wrong direction, and the corresponding temporal distance. We evaluate this cost function for all the possible ordered pairs (*𝑣*_*stop*_, *𝑣*_*start*_) in order to build a cost matrix of all possible one-to-one connections of root starts and root stops. This cost matrix is used to compute the optimal pairwise matching using the Hungarian algorithm (Kuhn, 1955). The algorithm works as follows:

~~~
Function: Iterative reconnection of isolated root start and dead-ends
---------------------------------------------------------------------
→ Compute the cost matrix C, which values are the connection cost of all possible pairs (root stop, root start)
→ While there is at least one root stop and at least one root start left, do :
  → Compute the optimal set S of connections using the Hungarian algorithm
  → In S, select the connection of minimum cost and build the corresponding edge *e* = (*𝑣*_*stop*_, *𝑣*_*start*_).
  →If ((*t*(*𝑣*_*stop*_) = *t*(*𝑣*_*start*_)) and (insertion of *e* create a cycle in *F*):
    → reject *e* and stop the algorithm.
  → Else Add the new edge *e* to the graph
  → In C, update the values that could have been modified by adding *a*.
end
~~~

While the Hungarian algorithm provides a global and optimal set of connections, we do not apply this set in one go. We iterate over these steps: computation of the Hungarian algorithm, connection of the pair of minimal cost, and update of the cost matrix. That way, each new connection adds knowledge to the system. This point is critical for tackling some challenging cases, like when a single root is reconstructed from a series of numerous small isolated elements comprising both a root start and a root end. This kind of situation happens for example, when a lateral root grows hidden below a primary root (see results, Figure 6-D).

**Figure 4:**
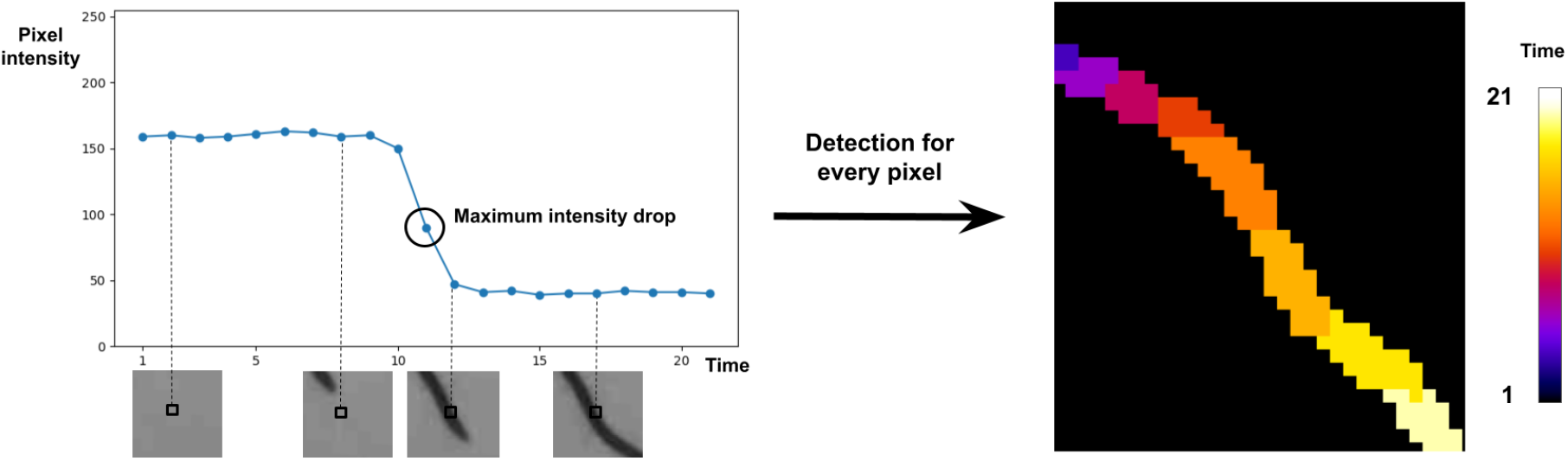
Principles of 2D+t segmentation. On the left: Intensity drop of a pixel (black square) when a root traverses it during the experiment and detection of the corresponding observation time (black circle). On the right: output of this operation when applied to a single growing root.

**Figure 5:**
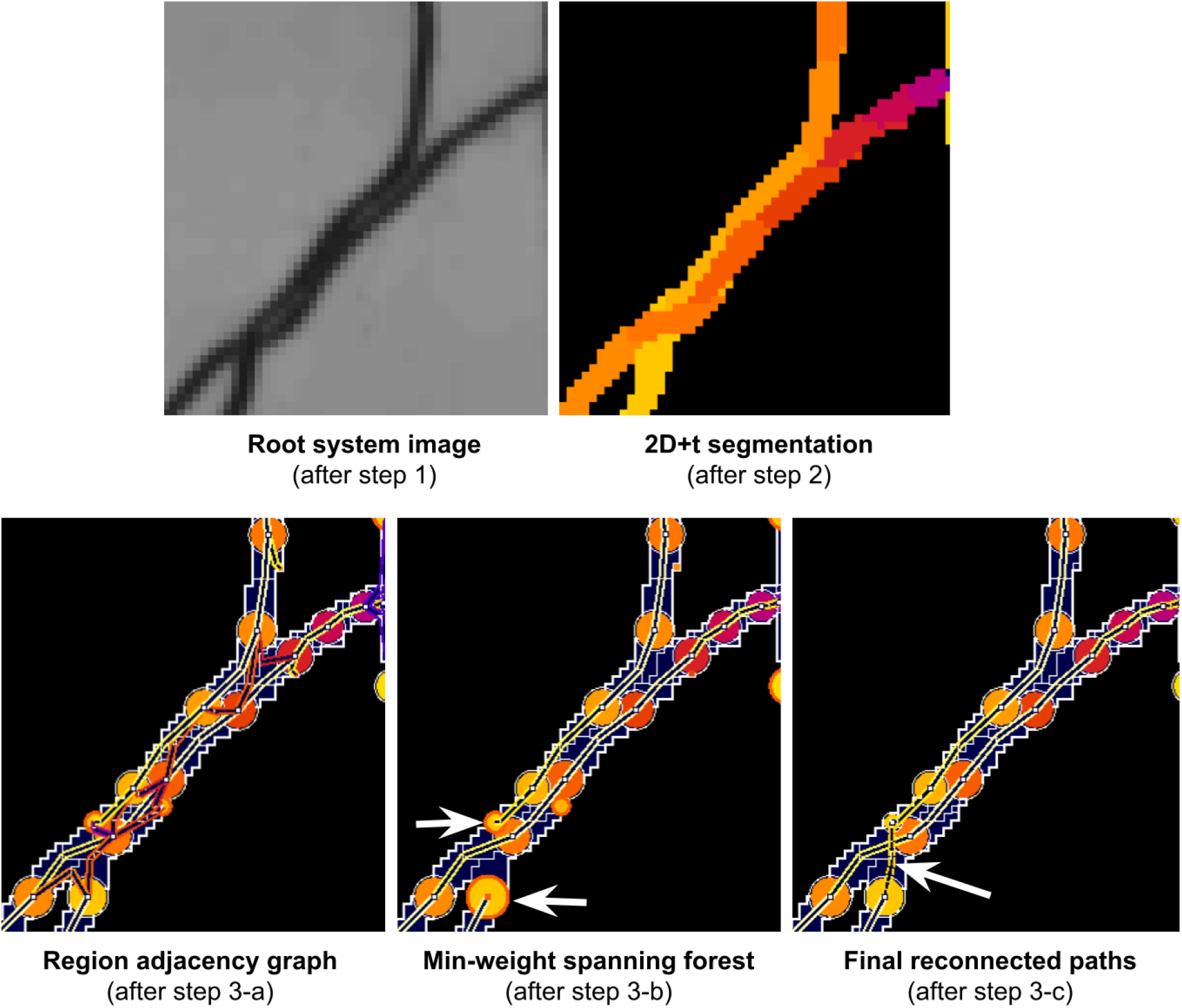
Successive operations during step 3. Upper row: output of steps 1 and 2. Bottom row; from left to right: construction of the initial graph (3-a), extraction of min-weight spanning forest (3-b), and root crossing resolution (3-c). The vertices colormap indicates their label, from 4 (red) to 13 (yellow), and the colormap of edges indicates their weight, from 0 (bright yellow) to 2 (red). White arrows: in the case of root crossing, the corresponding pathways in the graph are disconnected during step 3-b and reconnected in step 3-c.

**Figure 5:**
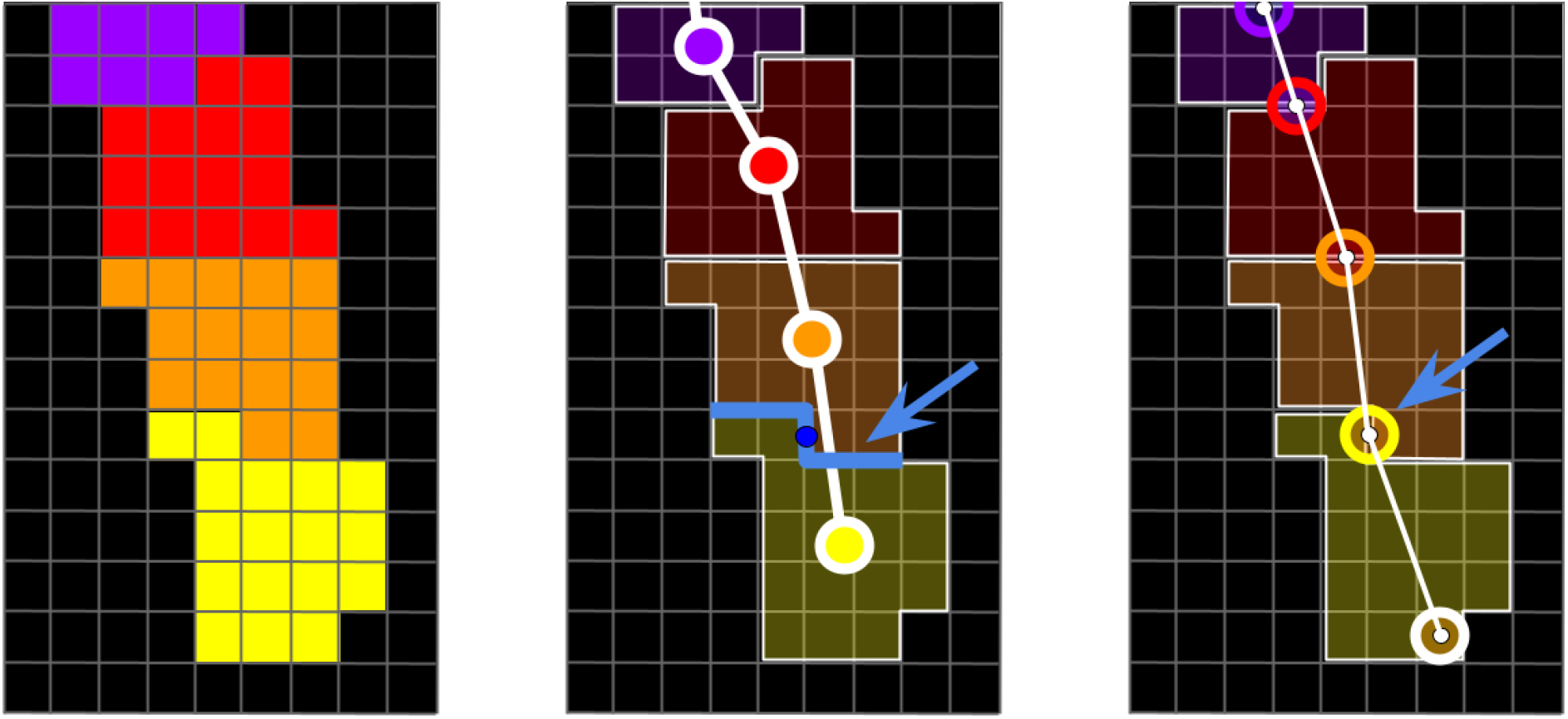
Estimation of root central line 2D+t coordinates. Left: labeled segmentation, comprising successive growth intervals of a single root, from purple (earliest interval) to yellow (latest interval). Middle: corresponding elements (edges and vertices) in the min-weighted spanning forest F extracted from the region adjacency graph. Right: Corresponding elements in ℑ, the dual graph of F, whose spatial coordinates of vertices are estimated from the geometry of the separation between successive growth intervals (blue arrow and blue line).

**Figure 6:**
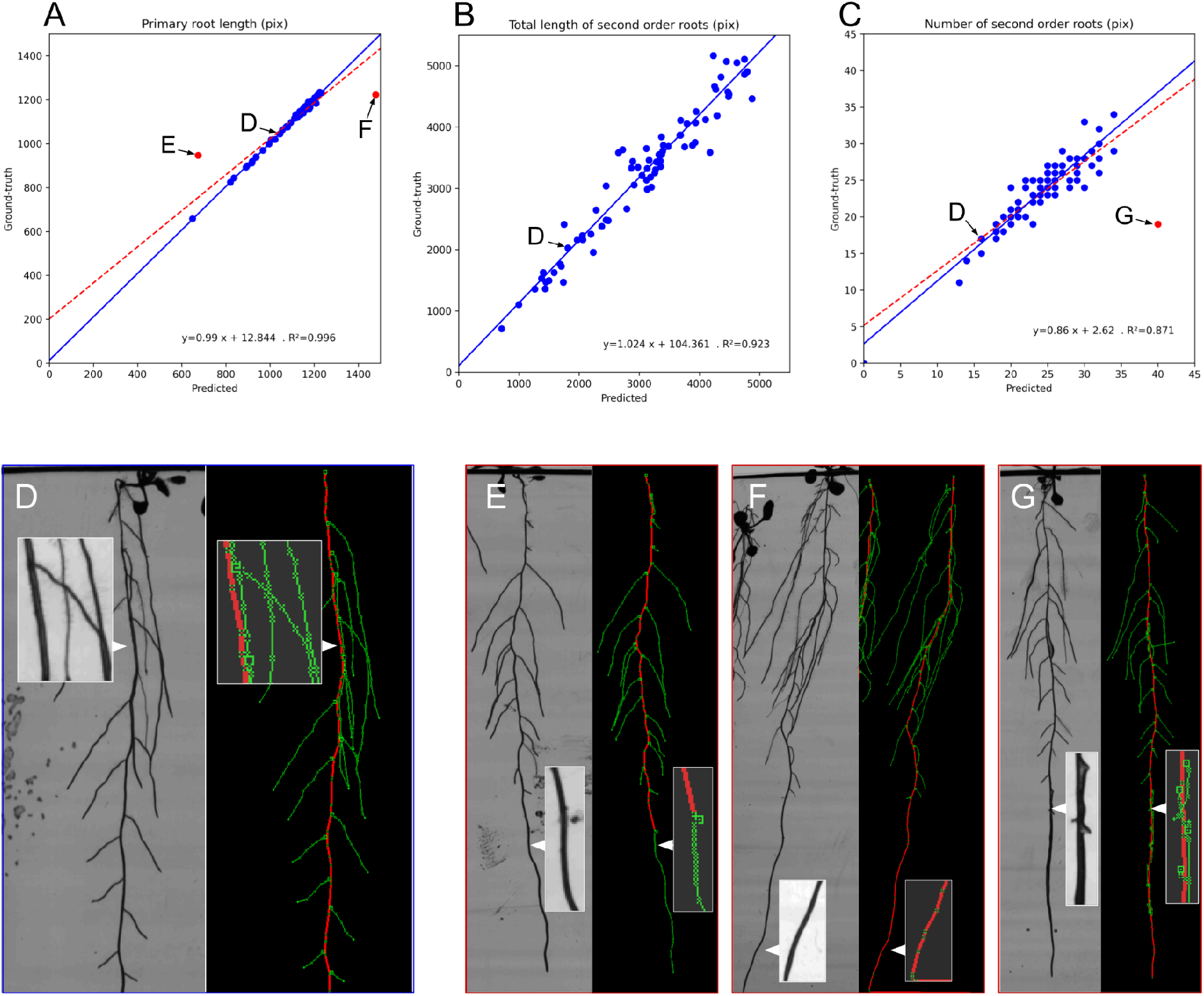
Performances in the estimation of static traits by comparison between predicted and expertized data. Traits investigated are: A) primary root length, B) total second-order root length, and C) number of second-order roots. The three red points in A, B, and C, are outliers that were not considered for linear regression (in blue). The four images, D, E, F, and G, present the source image and its associated reconstructed RSA. Each corresponds to a point in graphs A, B, and C. In D), the reconstruction process captured the architecture efficiently despite many ambiguities: numerous root crossings and a second-order root following the trajectory of the primary root. The three outliers E), F), and G) occurred in unexpected situations leading to significant reconstruction errors. In E), an active bacterial strain is present on the path of the primary root, yielding a false positive of the primary root detection at an early stage and the interruption of the primary axis path. In F), the root reaches the bottom of the petri dish. While this common situation generally does not impact the reconstruction, in this case, a dark structure at the bottom of the dish is interpreted as the subsequent path of the primary root. In G), the primary root is infected with a bacterial strain. As the bacterial strain develops, the root exhibits a rapid increase in apparent thickness, interpreted as the emergence of many lateral roots.

### 3-d) Artifacts rejection

We compute statistics over the identified lateral root paths and proceed to detection of outliers in order to reject spurious lateral roots. To that end, we collect data characterizing the geometry and dynamics of these paths, measured at each timestep: length, speed, and surface. These scalar values are binned with respect to the temporal distance from the emergence of the path. Then we compute robust statistics within these subgroups. We compute the double-sided MADe (Leys et al. 2013) and detect values exceeding twenty-five times the median absolute deviation around the median. These abnormal values are identified as markers of spurious roots. The corresponding roots are then rejected as artifacts.

### 4 Architecture reconstruction

The reconstruction of the spatiotemporal model of the RSA involves a precise description of the topology and geometry of the root systems. Vertices and edges of the forest *F* resulting from step 3 describe the root system topology. However, they do not provide the precise spatiotemporal pathway: each vertex of *F* represents a growing region of root pixels between two observations; thus, it is not associated with any exact coordinates, neither in time nor space. To build a precise spatiotemporal model, we need to know the precise position of each root tip at each observation time point and then interpolate this position between the time points to follow the roots’ actual pathway, whatever their speed and tortuosity.

The position of the root tip at each time point can be estimated from the edges of the forest *F*, recalling that each edge *e* ∈ *F* corresponds to a transition between two consecutive growth areas of a root (see figure 5). Thus, edges of *F* correspond to the successive positions of root tips at each time point. The spatiotemporal model is then represented by the dual graph ℑ = (*V*_ℑ_, *E*_ℑ_) of *F* = (*V*_*F*_, *E*_*F*_) where a vertex encodes the spatiotemporal position of a root tip and an edge encodes the root segment between two positions (see Figure 5). In this dual graph (Figure 5, right), a vertex *w* ∈ ℑ corresponds to an edge in *F*. Its temporal coordinate is well-defined; it is exactly an observation time point.

Conversely, the spatial coordinates of *w* ∈ ℑ have to be defined. *w* corresponds to an adjacency relationship between two distinct connected components in *S* (see the blue line and the blue arrow in Figure 4, center). We define the geometry of this adjacency in the image space by extracting the set of pixel edges and corners separating the two adjacent components. We choose the spatial coordinates of *w* ∈ ℑ to be the coordinates of the central-most pixel corner or pixel edge over all the corresponding frontiers (see the dark blue circle in Figure 4, center). We process all the edges of *F* that way to compute the corresponding vertices of ℑ. Afterward, we create for each root an additional vertex *w* ∈ ℑ to identify the root tip. We estimate its position by identifying the pixel of the last component that is the most distant (geodesically) from the nearest identified vertex *w* ∈ ℑ (for an illustration, see the lower white circle in Figure 4, right). Once all the vertices of ℑ are computed this way, we connect them with edges corresponding to the vertices of *F*. The geometry of the edges is defined by straight lines connecting successive positions of root tips. These edges constitute an oversimplified version of the root curves, which do not precisely follow the roots pathway, especially for primary roots, which are growing very fast. We refine ℑ to ensure geometrical fidelity of the RSA model, allowing accurate measurements (especially root length). For each edge ε = (*w*_1_, ω_2_)∈ ℑ, we compute the shortest path between *w* and *w* in the corresponding connected component *CC* ∈ *S*. We use the Dijkstra algorithm and compute the shortest path of 8-connected pixels in *CC*. We define the edge cost using the standard inter-pixel euclidean distance, weighted by a scalar function derived from the distance map of the traversed pixels to the contour of the *CC*. This morphological feature forces the path to avoid understeering curves; it fosters representative RSA models.

Finally, we reduce the number of points in the pixel path by decimating it using the Douglas-Peucker algorithm (Douglas and Peucker, 1973). The temporal coordinate of the vertices is estimated by a linear interpolation computed from the nearest two bounding vertices whose temporal coordinates are known.

### Upstream extrapolation of primary roots to the seed

Leaves sometimes falls into the medium during the time course, inducing false detection of new organs. We address this problem by artificially removing the falling leaves from the images using a series of morphological operations. However, this results in a loss of information about the primary and lateral root starts, causing the pipeline to detect the root system initiation point below the target point (the seed). The pipeline corrects this defect during this post-processing operation by computing a pixel path of minimum cost from the target height to the detected initiation point using the Dijkstra algorithm. To that end, we define the cost of pixel-to-pixel traversal as the product of the inter-pixel distance (1 for pixel sharing an edge or 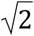 for pixel sharing a corner) by the average intensity of the two pixels.

### Downstream extrapolation of the stopped laterals

After computation, we check every root end, and investigate the one ending before the last observation time, probably corresponding to lateral roots stopping their growth before the sequence ending. Such an event is infrequent, thus suspect. If these early-stopping roots *r*_*stopped*_ are not incident to any other root, we consider that *r*_*stopped*_ have stopped its growth. In that case, we keep it in the model if it had a continuous growth during at least three time points, or we reject it instead. In the other case where *r*_*stopped*_ is incident to another earlier root *r*_*first*_, we assume that *r*_*stopped*_ continued its growth hidden below or parallel to *r*_*first*_. In this situation, we extrapolate the growth of *r*_*stopped*_ to its estimated final position in the time-sequence and add these new points to the RSA model.

## Results

We built an annotated dataset for validation purposes. The performance of the pipeline is measured using several metrics to estimate the geometrical and topological fidelity of the reconstruction.

### Pipeline validation

#### Validation dataset

The entire dataset was acquired within a seven days period. It contained *N*_*tot*_ = 1000 plants, divided into 200 petri dishes loaded into HIRROS. Experiments took place over 160 hours, with one observation every 8 hours (each plant/dish is observed 21 times): the total image dataset contains 21.000 plant.timestep. We selected a representative subset of 36 Petri dishes (*N*_*test*_ = 180 plants) to be manually annotated along the 21 time points (3780 plant.timestep) to perform validation tests. These 36 dishes cover the genotypic and environmental variations. The first 18 Petri dishes are used to train the model and set the parameters.

The 18 remaining dishes are used to compute the validation metrics. The results presented hereafter are computed on a test set of 18 × 5 × 21 = 1890 plant.timestep.

### Validation methodology

Assessing the pipeline’s performance involves confronting the pipeline’s outputs with corresponding data annotated by an expert to assess the quality of the analysis and identify drawbacks. We used the Smartroot software library (Lobet *et al*., *2011)* to develop an ImageJ plugin to edit the pipeline’s output. Our plugin provides a simple GUI to navigate an RSA model in 2D + t and apply elementary editing operations: adding, modifying, and removing a root or a vertex. Each edition operation takes place in 2D + t. After each operation, the topology is checked automatically to correct the graph if needed. The expert stops editing the RSA model when he estimates that both the topology and geometry are valid at each time point of the sequence of observations. In our dataset, the mean number of operations to correct the RSA of a single plant was 15.4 +- 6.3 operations per plant (mean +- standard deviation).

The expert identifies which plant can be considered an outlier during the annotation process. Most of the time, outliers are bacteria-contaminated plants that stopped growing or died during the time course. These plants are removed from the dataset and are not included in the validation dataset.

### Performance of static trait estimation

We first assess the performance of the pipeline on the estimation of static traits by using global traits frequently used in the literature: total predicted number of organs and predicted cumulated length for primary and second-order roots. Our dataset is composed of complex root architectures with numerous organ crossings. We compute these traits at the last time point when the architecture reaches its maximum complexity. At this stage, we evaluate the capabilities of our method to disentangle the numerous organs when processing complex root systems.

The three graphs in figure 6 illustrate the accuracy of the measurements, especially on organ length estimation (figure 6A and B), with a correlation coefficient of 0.996 and 0.923 for primary and lateral roots, respectively. Moreover, regression lines between predicted and ground-truth values have small intercepts and slope coefficients near 1 (i.e., 0.99 for primaries and 1. 02 for second-order roots, see Figures 6-A and 6-B). Our method, applied on a complex dataset, also exhibits state-of-the-art performances (Yasrab et al. 2019; Yasrab et al. 2021) on second-order root counting (figure 6C), with a correlation coefficient of 0.818. These performances make our method suited for collecting the most common phenes required in high-throughput experiments. Figures 6E, 6F, and 6G highlight three outliers (∼ 3% of the dataset) due to developing bacterial strains. These non-plant elements were identified as roots and severely impacted the prediction of the pipeline.

### Performance of dynamic traits estimation

The pipeline reconstructs RSA over time, allowing to quantify each root’s elongation rate and its evolution throughout the time series, providing access to dynamic traits. In this section, we evaluate the pipeline performance of dynamic traits quantification. We measured the total additional length between successive time points and the additional length corresponding to laterals. We further investigate the source of errors by estimating the additional length of laterals within three classes: roots attached to the primary’s upper part, middle part, and the bottom part (Figure 7).

**Figure 7:**
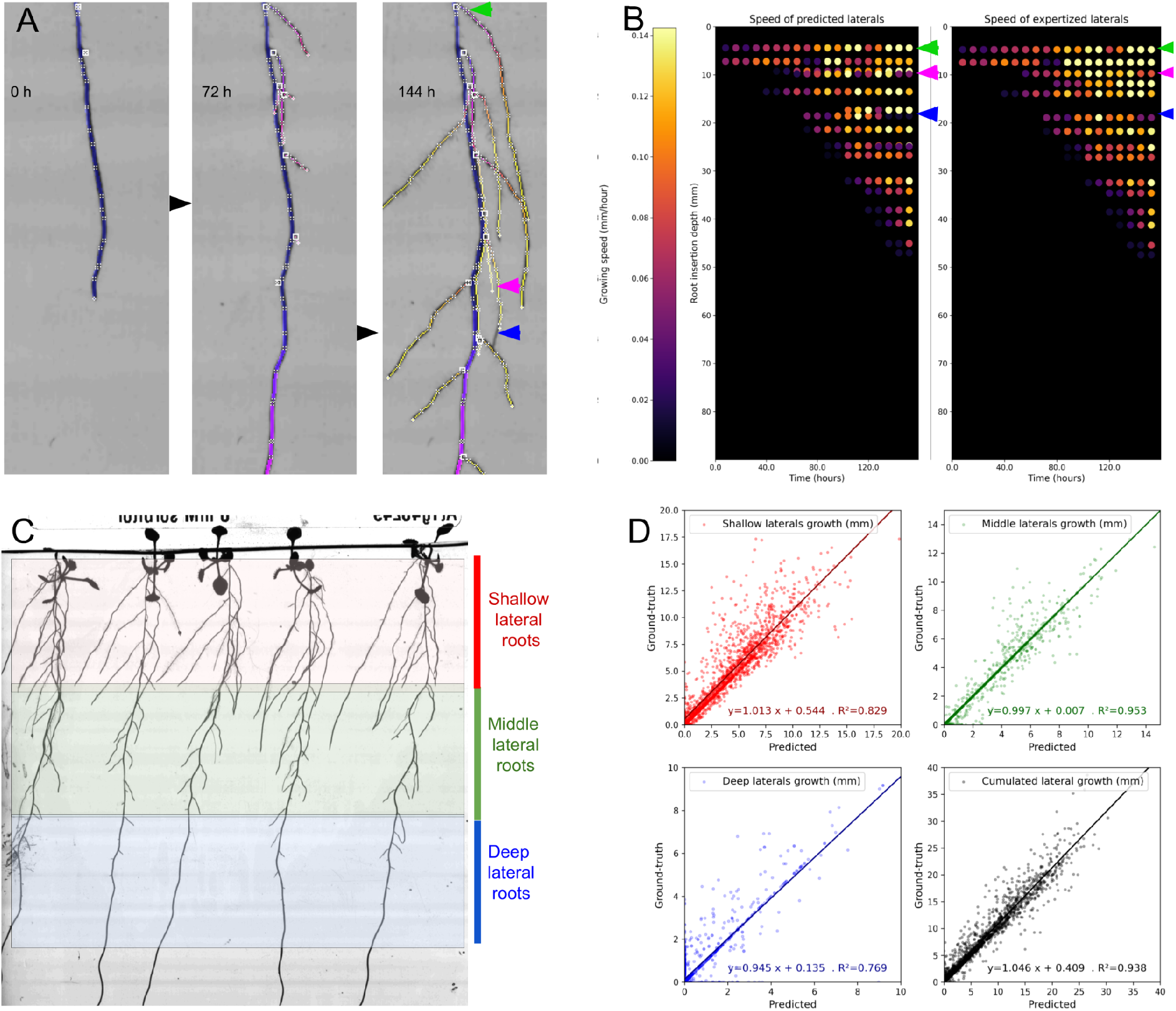
Performance of quantifying dynamic traits of lateral roots emergence and elongation rate by comparing predicted and expertized data. **A)** 2D+t reconstructed RSA. **B)** Predicted individual lateral roots growth speed (left) vs. annotated data (right). **D)** Performance of estimating the total lateral root growth of individual plants between two consecutive observations (in black). The performance in estimating individual root stages growth (shallow, middle, and deep root stage growth, see **C)**) are shown respectively in red, green, and blue.

Figure 7A represents the successive observation of a root system overlaid with the corresponding reconstructed architecture. Figure 7B represents the 2D+t reconstruction of a unique plant from data automatically obtained by the pipeline (left panel) or from expert analysis (right panel). Each horizontal dot-line represents a lateral root, and the color of each dot represents the elongation rate of the individual root between 2 successive time points. This figure visually illustrates the high accuracy of the analysis. Most organs are well estimated (green arrow), with minor differences occurring when the track of a root is lost and prolonged artificially (pink arrow) or when a root is cut into two single organs (blue arrow). The cumulated lateral root growth is estimated between two successive observations with high accuracy (*R*^2^ = 0. 938). Regression lines between predicted and ground truth values have small intercepts and a slope coefficient near 1 (i.e., 1.05). The most challenging part to estimate is the deeper compartment of lateral roots, inserted in the deepest third of the primary axis. Roots growth from this compartment is estimated with a lower precision (*R*^2^ = 0. 769, regression line slope = 0. 95). These roots appear at the end of the time course, are smaller, and the length of the growing units is closer to the pixel size. Thus their growing speed cannot be estimated accurately. The shallowest third (Figure 6-C, in red) suffers from other misestimation biases induced by reconstruction errors when the leaves fall into the root system (*R*^2^ = 0. 829, slope = 1. 01). Finally, the middle compartment roots do not suffer from these issues and exhibit a high accuracy (*R*^2^ = 0. 953, slope = 1. 00). To our knowledge, there are no phenotyping studies that validate such dynamic traits, while these measurements are of high interest to plant scientists.

### Characterization of spatiotemporal phenotypes of root systems

In this section, we investigate the resulting spatiotemporal traits extracted to unravel the studied root systems’ characteristics and assess our pipeline’s interest for GxE studies.

#### Topological and geometrical characterization of root system dynamics

Root system topology is widely complex, requiring the information to be simplified before being analyzed. We derived an approach used for meristem organ characterization to produce a heatmap of the growing areas of the root systems. In order to evaluate this approach, we compared the reconstructed graph of plants grown for six days on a control medium or subjected to a 150 mM sorbitol osmotic stress (Figure 8). Terminal RSA analysis showed that osmotic stress reduces primary and lateral root length by 16 and 50%, respectively (Table 1). Kinetic reconstruction clearly showed that the primary root elongation rate was reduced under osmotic stress since the application of treatment (green line) but appeared more variable between plants than under control conditions (dark green cone). On the contrary, osmotic stress dramatically delayed the development of older lateral roots in the 0-15 mm depth interval and significantly reduced the elongation rate of these roots (colder color on the heat map). On the contrary, lateral roots on the 30-50 mm depth interval appeared less affected by osmotic stress with no significant delay in development and little difference in the elongation rate, suggesting different sensitivity or acclimation to osmotic stress. Finally, osmotic stress also altered the topology of the root system by reducing the distance from the last visible lateral root to the primary root tip (distance between the green line and the last visible line of the heat map). Altogether, these results open promising and unexpected perspectives to analyze the dynamics of the RSA response to osmotic stress.

**Table 1:** Comparison of primary and lateral root length of two populations with different GxE conditions

|  | Primary root length | Lateral root length |
| --- | --- | --- |
| Control (mean +- std over N=30) | 80.6 +- 8.5 | 184.7 +- 36.8 |
| Sorbitol, 150 mM (mean +- std over N=30) | 69.2 +- 15.0 | 89.7 +- 41.3 |

**Figure 8:**
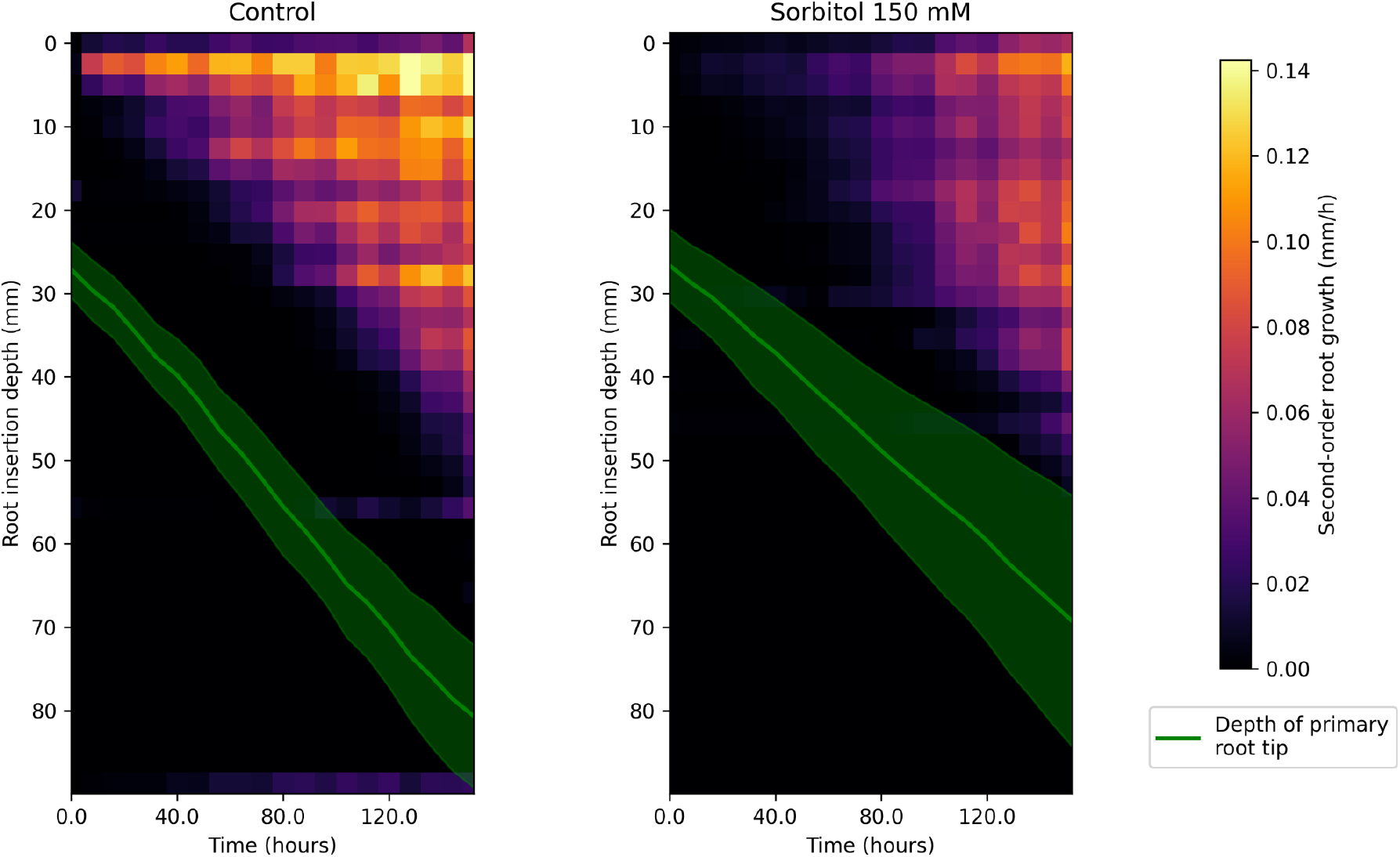
Elongation rates of second-order roots as a function of their insertion depth on the primary root. Each graph results from statistics computed over N=30 specimens of the same genotype grown under control conditions or subjected to osmotic stress (150mM sorbitol). Primary root tip depth is displayed in green (mean +- std-dev).

**Figure 9:**
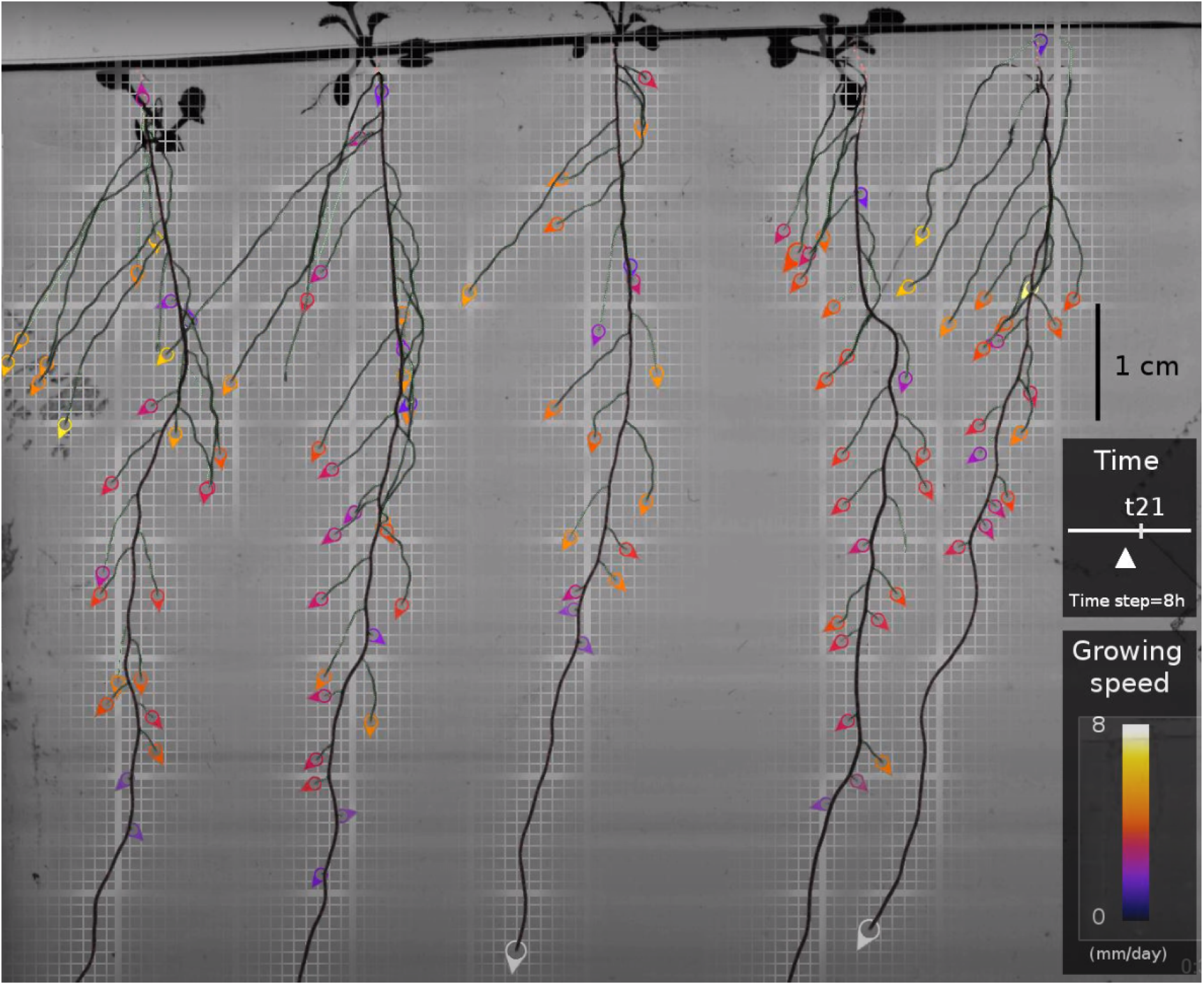
Screenshot of the example movie. Accessible at https://www.youtube.com/watch?v=SWEqnnOhIOU

### Time-lapse visualization of the growing root system

In addition to the RSA reconstruction pipeline, we propose a method to visualize the growth of the root system. The resulting movie shows the compilation of a 7-days growth in one minute. It gathers all the pipeline achievements in a realistic rendering, giving a rapid and accurate overview of the reconstruction of the experiment and facilitating investigations by non-computer specialists.

The movie is built using multiple interpolation techniques. First, the reconstructed architecture is rasterized in the pixel space. The apparition time of each pixel of the root central lines is determined by interpolating the spatiotemporal coordinates of the surrounding vertices in the model. Each pixel of the segmented RSA is then given the apparition time of the nearest pixel of the corresponding central line. After this operation, the binary segmentation of the whole architecture is parameterized by a time function, giving the apparition time of each pixel. This 2D+t segmentation is used to interpolate successive masks of the growing RSA that are applied on the original keyframes 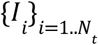 of the registered 2D time series.

The root system mask obtained for a time *T*_*i*_ < *t* < *T*_*i*+1_ is applied to the image *I*_*i*+1_ to build the root architecture part while its complementary is applied to the image *I*_*i*_ to gather the background part. The assembling is done by summing these two images.

When *t* is close to an observation time *T*_*i*_ (images near a keyframe), we proceed to a fading by computing the same operations using the subsequent keyframes (*I*_*i*+1_ and *I*_*i*+2_). Finally the individual results are merged with fading factors that vary following a linear ramp to make these transitions unnoticeable.

After interpolating time-lapse images, we compute the position and speed at each time frame from the reconstructed RSA. They are overlaid on the image and displayed as colored tips with a colormap indicating the elongation rate.

The default parameters of the procedure result in a movie of the growing RSA generating one image every 15 mn, from an initial time series composed of 21 timeframes, one every 8 hours.

An example movie is available online at this youtube link: https://www.youtube.com/watch?v=SWEqnnOhIOU. The provided open-source software can generate such a movie from any of our 200 reconstructed time sequences in approximately one minute.

## Discussion

In this contribution, we proposed a novel method for robust acquisition of root system architecture with its topology and development with HIRROS, a high-throughput root phenotyping platform. Our automatic pipeline combines spatial and temporal information to segment structures with their apparition time and a topological tracking algorithm to reconstruct the whole architecture, including primary and second-order roots. Our pipeline is robust to root crossing. We evaluated our method on a dataset of complex root systems and assessed performance in terms of geometric and topologic fidelity of the reconstructed 2D+t architectures. We show the ability of our method to capture spatio-temporal phenes of interest to the community, with a high level of guarantees.

We developed a high-resolution automated imaging system that can handle up to 200 Petri dishes (i.e., up to 1000 plants) and image the plates at high resolution (19 µm per pixel) and high contrast allowing excellent root/background separation. Using a telecentric lens avoids image distortion and provides a huge depth of field combined with a micrometric controlled plate translation. HIRROS combines the high-resolution imaging as proposed by Gaggion *et al*., 2021 and Ohlsson *et al*., 2021, with high-throughput (Nagel et al., 2020), offering a unique opportunity to explore the GxE root adaptive responses to a wide range of abiotic stresses. Petri dishes that contain transparent agar gelified medium are commonly used for *Arabidopsis thaliana* and model plants RSA analysis. However, they imply several constraints like root crossing, secondary roots juxtapositions, and shoots’ movement due to 2D space limitation. These physiological and technical constraints restrict RSA analysis to static or global dynamic parameters (Gaggion *et al*., 2021). In order to access the dynamic responses of the individual roots, we developed a dedicated image analysis pipeline.

Our proposition differs from most state-of-the-art solutions, where segmentation is performed separately on successive 2D images, which involves a subsequent step to match the successive segmentations or RSA reconstructions (Yasrab et al., 2021; Möller et al., 2021). One solution is to use a well-trained neural network to perform root segmentation successfully in each 2D image using the information from the previous images (Yasrab et al., 2021). However, the training this type of network requires a large training set of manually annotated images and a tedious task of reannotation in case of transfer learning. We propose a novel method incorporating a registration step to achieve accurate superposition of successive observed images and avoid tree matching problems. The registration step produces a 2D+t stack of aligned images, which enables us to propose a pixel-wise root detector that performs root segmentation and determination of root pixel apparition time in one go. Our pipeline combines spatial and temporal information, facilitating the segmentation step to such an extent that our mean-shift detector achieves performance comparable to deep neural networks (Yasrab et al., 2021, Möller et al., 2021).

However, our method has shown some limitations. When unwanted structures appear in the image field, they can be mistaken for growing roots, resulting in errors that must be detected and post-processed during the graph construction step. This point complicates the algorithm, which must adapt to specific circumstances by deploying additional post-processing operations. The pipeline could benefit from a pre-processing step integrating a neural network in charge of predicting the pixel-wise probability of the presence of a root and removing unwanted structures from the input of the mean-shift detector. That way, we could take the best of the two worlds and probably achieve even better performances, although this prospect implies that our pipeline must be trainable. As such, our pipeline does not require any tedious annotation and training tasks.

Nevertheless, while our pipeline is fully automatic, some parameters need to be calibrated for a new experimental configuration: mean intensity of roots, mean intensity of background, and max acceptable root speed. In a trainable version of the pipeline, these parameters could be derived automatically from a dataset containing some annotated examples. By integrating these improvements, our pipeline could be transferable to other datasets.

Analysis of root system growth in Petri dishes is a unique tool for high-throughput phenotyping. However, observing root systems in 2D involves dealing with multiple root crossovers as root system complexity increases. These situations are difficult to disentangle and can hinder the tracking of individual roots. Many algorithms have been proposed to address this challenge. Most of them are greedy algorithms, processing root after root or time after time (Lobet et al., 2011; Pound et al., 2013; Möller et al., 2021; Yasrab et al., 2021) without a global view. In this work, we design a global tracking algorithm based on the region adjacency graph extracted from the segmentation of individual root segments. We evaluated the success of this algorithm in challenging cases when laterals hide below primaries or multiple laterals (N=3, 4, 5+) intersect in a small space.

The formal framework of our algorithm relies on strong assumptions: there is no third-order ramification, and through time all the growing roots pixels are included in the final image. Our results show that some specimens do not meet these assumptions, which induces reconstruction errors, such as when plants are infected with bacterial strains. Moreover, our algorithm is designed to accurately track one primary root per plant and its second-order roots. It does not include the possibility of multiple primary roots or third-order organs. Further work will focus on adjusting the assumptions to deal with the emergence of third-order organs and to work with other species with different root system architecture patterns.

Performances of the method have been evaluated using standard metrics on a complex architecture dataset. Interestingly, the relative error of the total lateral length does not seem to increase with the development of the root system. This point could be crucial to study more complex architectures. In addition, we validated the pipeline’s ability to estimate organs’ growth rate. By providing such guarantees, we open the possibility of using these dynamic phenes in GxE studies. We illustrated this point by comparing plant development under various levels of osmotic stresses, highlighting differences in primary and lateral growth patterns. Finally, we developed a visualization method to construct a simulated time-lapse growth movie. This method opens up the possibility of generating a large amount of realistic synthetic data with a parameterizable timestep and voxel size, leading to the design of a trainable model.

## Conclusion

The presented method provides a unique solution for high-throughput phenotyping of root system architecture. It is robust to root crossing, which is the main difficulty for accurate tracking of the individual organs from 2D image time-series. The reconstructed architecture is provided in RSML format, which integrates the topology and geometry of the root system, parameterized with the appearance time of the root segments.

Further work will focus on validating this pipeline with a larger panel of GxE variations and longer times series, adapting the hypothesis used for the topological tracking, and making the pipeline trainable to transfer it to different species or mounting configurations (box size, rhizotron). The resulting architectures will be used as topological and geometrical models to feed functional and structural root models and further understand root system development.

## Software availability

The architecture reconstruction pipeline is supplied as an ImageJ plugin with online documentation (Plugin page: https://imagej.net/plugins/rootsystemtracker [Fernandez, 2022]) and as open-source code on GitHub (https://github.com/Rocsg/RootSystemTracker).

Once the reconstruction is achieved, the architecture can be investigated using the 2D+t model edition plugin, can be used to extract other phenes using RSML reader modules, or can be used to reconstruct a time-lapse movie.

## References

Balduzzi M, Binder BM, Bucksch A, Chang C, Hong L, Iyer-Pascuzzi AS, Pradal C, Sparks EE. Reshaping plant biology: qualitative and quantitative descriptors for plant morphology. Frontiers in Plant Science. 2017 Feb 3;8:117.

Boudon F, Preuksakarn C, Ferraro P, Diener J, Nacry P, Nikinmaa E, Godin C. Quantitative assessment of automatic reconstructions of branching systems obtained from laser scanning. Annals of Botany. 2014 Sep 1;114(4):853–62.

Boursiac Y, Pradal C, Bauget F, Lucas M, Delivorias S, Godin C, Maurel C. Phenotyping and modeling of root hydraulic architecture reveal critical determinants of axial water transport.

Delory BM, Li M, Topp CN, Lobet G. archiDART v3. 0: A new data analysis pipeline allowing the topological analysis of plant root systems. F1000Research. 2018;7.

Douglas DH, Peucker TK. Algorithms for the reduction of the number of points required to represent a digitized line or its caricature. Cartographica: the international journal for geographic information and geovisualization. 1973 Dec 1;10(2):112–22.

Edmonds J. Optimum branchings. Journal of Research of the national Bureau of Standards B. 1967 Oct;71(4):233–40.

Fernandez R, Moisy C. Fijiyama: a registration tool for 3D multimodal time-lapse imaging. Bioinformatics. 2021 Jun 16;37(10):1482–4.

Fernandez R. RootSystemTracker plugin page https://imagej.net/plugins/rootsystemtracker. Accessed 12 Jul 2022

Gaggion N, Ariel F, Daric V, Lambert É, Legendre S, Roulé T, Camoirano A, Milone DH, Crespi M, Blein T, Ferrante E. ChronoRoot: High-throughput phenotyping by deep segmentation networks reveals novel temporal parameters of plant root system architecture. GigaScience. 2021 Jul;10(7):giab052.

Griffiths M, Mellor N, Sturrock CJ, Atkinson BS, Johnson J, Mairhofer S, York LM, Atkinson JA, Soltaninejad M, Foulkes JF, Pound MP. X-ray CT reveals 4D root system development and lateral root responses to nitrate in soil. The Plant Phenome Journal. 2022;5(1):e20036.

Herrero-Huerta M, Meline V, Iyer-Pascuzzi AS, Souza AM, Tuinstra MR, Yang Y. 4D Structural root architecture modeling from digital twins by X-Ray Computed Tomography. Plant methods. 2021 Dec;17(1):1–2.

Kuhn HW. The Hungarian method for the assignment problem. Naval research logistics quarterly. 1955 Mar;2(1-2):83–97.

LaRue T, Lindner H, Srinivas A, Exposito-Alonso M, Lobet G, Dinneny JR. Uncovering natural variation in root system architecture and growth dynamics using a robotics-assisted phenomics platform. Biorxiv. 2021 Jan 1.

Leys C, Ley C, Klein O, Bernard P, Licata L. Detecting outliers: Do not use standard deviation around the mean, use absolute deviation around the median. Journal of experimental social psychology. 2013 Jul 1;49(4):764–6.

Lobet G, Pagès L, Draye X. A novel image-analysis toolbox enabling quantitative analysis of root system architecture. Plant physiology. 2011 Sep;157(1):29–39.

Lobet G, Pound MP, Diener J, Pradal C, Draye X, Godin C, Javaux M, Leitner D, Meunier F, Nacry P, Pridmore TP. Root system markup language: toward a unified root architecture description language. Plant physiology. 2015 Mar;167(3):617–27.

Luo W, Xing J, Milan A, Zhang X, Liu W, Kim TK. Multiple object tracking: A literature review. Artificial Intelligence. 2021 Apr 1;293:103448.

Lynch JP. Harnessing root architecture to address global challenges. The Plant Journal. 2022 Jan;109(2):415–31.

Mairhofer S, Zappala S, Tracy SR, Sturrock C, Bennett M, Mooney SJ, Pridmore T. RooTrak: automated recovery of three-dimensional plant root architecture in soil from X-ray microcomputed tomography images using visual tracking. Plant physiology. 2012 Feb;158(2):561–9.

Mairhofer S, Johnson J, Sturrock CJ, Bennett MJ, Mooney SJ, Pridmore TP. Visual tracking for the recovery of multiple interacting plant root systems from X-ray μCT images. Machine Vision and Applications. 2016 Jul;27(5):721–34.

Maurel C, Nacry P. Root architecture and hydraulics converge for acclimation to changing water availability. Nature plants. 2020 Jul;6(7):744–9.

Möller B, Schreck B, Posch S. Analysis of Arabidopsis Root Images--Studies on CNNs and Skeleton-Based Root Topology. InProceedings of the IEEE/CVF International Conference on Computer Vision 2021 (pp. 1294–1302).

Nacry P. Root Phenotyping platform webpage. 2022. https://www1.montpellier.inra.fr/wp-inra/bpmp/en/platform/root-phenotyping-platform. Accessed 12 Jul 2022

Nagel KA, Lenz H, Kastenholz B, Gilmer F, Averesch A, Putz A, Heinz K, Fischbach A, Scharr H, Fiorani F, Walter A. The platform GrowScreen-Agar enables identification of phenotypic diversity in root and shoot growth traits of agar grown plants. Plant Methods. 2020 Dec;16(1):1–7.

Ndour A, Vadez V, Pradal C, Lucas M. Virtual plants need water too: functional-structural root system models in the context of drought tolerance breeding. Frontiers in plant science. 2017 Sep 26;8:1577.

Ohlsson JA, Leong JX, Elander PH, Dauphinee AN, Ballhaus F, Johansson J, Lommel M, Hofmann G, Betnér S, Sandgren M, Schumacher K. SPIRO–the automated Petri plate imaging platform designed by biologists, for biologists. bioRxiv. 2021 Jan 1.

Ourselin S, Roche A, Prima S, Ayache N. Block matching: A general framework to improve robustness of rigid registration of medical images. In International Conference on Medical Image Computing And Computer-Assisted Intervention 2000 Oct 11 (pp. 557–566). Springer, Berlin, Heidelberg.

Smith AG, Han E, Petersen J, Olsen NA, Giese C, Athmann M, Dresbøll DB, Thorup-Kristensen K. RootPainter: deep learning segmentation of biological images with corrective annotation. BioRxiv. 2020 Jan 1.

Symonova O, Topp CN, Edelsbrunner H. DynamicRoots: a software platform for the reconstruction and analysis of growing plant roots. PLoS One. 2015 Jun 1;10(6):e0127657.

Takahashi H, Pradal C. Root phenotyping: important and minimum information required for root modeling in crop plants. Breeding Science. 2021;71(1):109–16.

Yasrab R, Atkinson JA, Wells DM, French AP, Pridmore TP, Pound MP. RootNav 2.0: Deep learning for automatic navigation of complex plant root architectures. GigaScience. 2019 Nov;8(11):giz123.

Yasrab R, Zhang J, Smyth P, Pound MP. Predicting plant growth from time-series data using deep learning. Remote Sensing. 2021 Jan 20;13(3):331.

